# dRep: A tool for fast and accurate genome de-replication that enables tracking of microbial genotypes and improved genome recovery from metagenomes

**DOI:** 10.1101/108142

**Authors:** Matthew R. Olm, Christopher T. Brown, Brandon Brooks, Jillian F. Banfield

**Affiliations:** Department of Plant and Microbial Biology, University of California, Berkeley, CA, USA.; Department of Environmental Science, Policy, and Management, University of California, Berkeley, CA, USA; Department of Earth and Planetary Science, University of California, Berkeley, CA, USA.

## Abstract

The number of microbial genomes sequenced each year is expanding rapidly, in part due to genome-resolved metagenomic studies that routinely recover hundreds of draft-quality genomes. Rapid algorithms have been developed to comprehensively compare large genome sets, but they are not accurate with draft-quality genomes. Here we present dRep, a program that sequentially applies a fast, inaccurate estimation of genome distance and a slow but accurate measure of average nucleotide identity to reduce the computational time for pair-wise genome set comparisons by orders of magnitude. We demonstrate its use in a study where we separately assembled each metagenome from time series datasets. Groups of essentially identical genomes were identified with dRep, and the best genome from each set was selected. This resulted in recovery of significantly more and higher-quality genomes compared to the set recovered using the typical co-assembly method. Documentation is available at http://drep.readthedocs.io/en/master/ and source code is available at https://github.com/MrOlm/drep.

---

Genome-resolved metagenomics involves the recovery of genomes directly from environmental shotgun DNA sequence datasets (Tyson et al., 2004). Metagenomic analysis of related samples from the same ecosystem is often employed to investigate compositional stability and spatial or temporal variation. The approach can also reveal microbial co-occurrence patterns and identify factors or processes that control organism abundances. Analysis of sample series data is also important technically, as different abundance patterns across the sample series for different organisms provide valuable constraints for binning of assembled fragments into genomes (Sharon et al., 2013). In this process, reads from individual samples are mapped back to a collection of genomes that is often obtained by combining the reads from all samples and assembling them together (co-assembly) (Vineis et al., 2016; Bendall et al., 2016; Lee et al., 2016). However, co-assembly dramatically increases the dataset size and complexity, especially when multiple different strains of the same species are present across the sample series, and can result in fragmented assemblies (Sczyrba et al., 2017).

An alternative process is to map reads to a collection of genomes independently assembled from the individual samples (**Supp. Figure S1**). Independent assembly should generate more and higher quality genomes than the co-assembly based approach because the complexity of individual samples is lower than that of the combination of samples. The challenge that arises from independent assembly is that de-replication of the resulting genome set is required (Olm et al., 2017; Raveh-Sadka et al., 2015; Probst et al., 2016). In addition to identifying genomes that are the “same,” another important aspect of de-replication is identifying the highest quality genome in each replicate set. This is important to maximize the accuracy of metabolic predictions and other downstream analyses.

De-replication requires pair-wise genome comparisons, and thus the time required scales exponentially with an increasing number of genomes. Hundreds of thousands of CPU hours may be needed to de-replicate larger genome sets with robust algorithms (gANI) (Varghese et al., 2015). Mash, a recently developed algorithm that utilizes MinHash distance to estimate similarity between genomes, is an attractive alternative due to its incredibly fast speed (Ondov et al., 2016). However, we found that the accuracy of MASH decreases as the completeness of the compared genome bins decreases (**Figure 1A**). Thus, it cannot be used to de-replicate collections of partial genomes.

**Figure 1.**
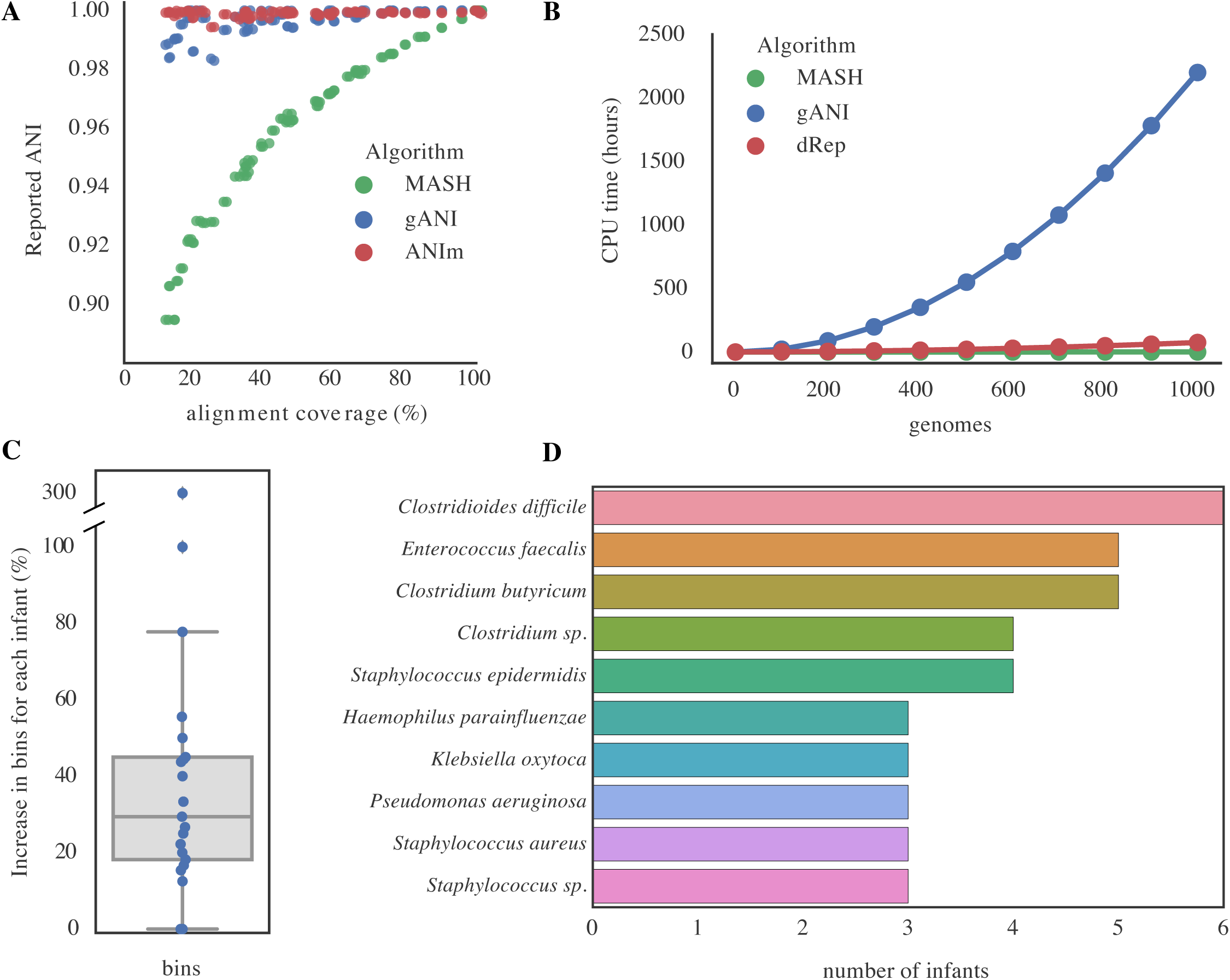
Assembly and de-replication with dRep results in more and higher-quality genome bins as compared to co-assembly. (**A**) A complete *Escherichia coli* genome was subset ten times in increments of 10% (10%, 20%, 30%, etc.). Subsets were compared to each other in a pair-wise manner (100 total comparisons) using three algorithms- ANIm, MASH, and gANI. For each pair of subsets, the percent overlap between the two genomes is shown on the x-axis (alignment coverage measured using mummer as implemented in ANIm), and the ANI reported from each algorithm is shown on the y-axis. ANIm and gANI are accurate when genomes overlap by ≥50%, but MASH is only accurate when genomes are essentially complete. (**B**) Using previously reported algorithm run-times, we estimated the time required to de-replicate genome sets of various sizes. gANI exhibits a sharp exponential climb, limiting its use on larger genome sets; MASH and dRep do not. (**C**) De-replication of bins from individual assemblies and a co-assembly (dRep assembly method) resulted in more bins (≥50% complete, ≤25% contaminated) than co-assembly alone. (**D**) Defining strains as having >99.9% ANI over 50% of their genomes, 10 strains are present in at least 3 of the 21 infants analyzed.

Here we present dRep, a program that utilizes both gANI and Mash in a bi-phasic approach to dramatically reduce the computational time required for genome de-replication, while ensuring high accuracy. The genome set is first divided into primary clusters using Mash, and then each primary cluster is compared in a pair-wise manner using gANI, forming secondary clusters of near-identical genomes that can be de-replicated. Using published information about time required for genome comparisons, we performed an *in silico* simulation of de-replication time for Mash, gANI, and dRep (Figure 1B). The results indicate that dRep affords a multiple orders of magnitude increase in computation efficiency compared to naïve gANI. To verify this prediction, we ran dRep on 1,125 genomes assembled from 195 fecal metagenomes collected from 21 premature infants during the first months of life (Raveh-Sadka et al., 2016), and found the actual run time to be very close to that predicted by our simulation: 92 versus 93 CPU hours, respectively. This is compared to 2,784 hours required for naïve gANI. As the run-time of dRep depends on the diversity of the genome set, and pre-term infant gut communities are especially non-diverse (Gibson et al., 2016), even greater increases in computational efficiency are expected from most other environments than predicted by our simulation.

We analyzed the same 195 metagenomes to test the prediction that, for each infant, individual assembly and de-replication would generate more and higher quality genomes than co-assembly of the read datasets. We de-replicated genomes obtained from assemblies generated from each sample individually as well as from a co-assembly (to recover low-abundance genomes), and recovered a genome set with 34% more bins (≥50% complete, ≤25% contaminated) than were obtained from co-assembly alone (Figure 1C). In cases where genomes were recovered using both methods, genomes assembled from individual samples were significantly less fragmented (N50; *p* = 9.6e-13) and more complete (*p* = 2.3e-8) (Wilcoxon signed-rank test). We also tested the ability of dRep to track strains present in multiple infants. We defined strains as the “same” if at least 50% of the genomes aligned with 99.9% average nucleotide identity, and identified 10 strains present in at least three of the infants (Figure 1D). Taken together, dRep enabled recovery of more and better genomes than co-assembly alone, and served as an effective tool for strain tracking.

To explore the effect of strain heterogeneity on assembly and genome recovery, we performed co-assemblies and individual dataset assemblies followed by dRep on a sample series from a single infant. The infant dataset has known strain heterogeneity (Sharon et al., 2013) and has been used for benchmarking many new tools (Luo et al., 2015; Eren et al., 2015; Graham et al., 2016). In the case of *Staphylococus hominis*, co-assembly generated a contaminated bin (i.e., many duplicate and triplicate single copy genes) (Figure 2A). In contrast, a near-complete, uncontaminated genome was recovered from several individual time-points. Previous work on the same dataset (Eren et al., 2015) has shown manual bin curation of the co-assembled bin with anvi’o can increase the *S. hominis* bin quality (73% complete; 6.6% redundant), but still not to the level of the un-curated bin from the individual assembly (98% complete; 0% redundant).

**Figure 2:**
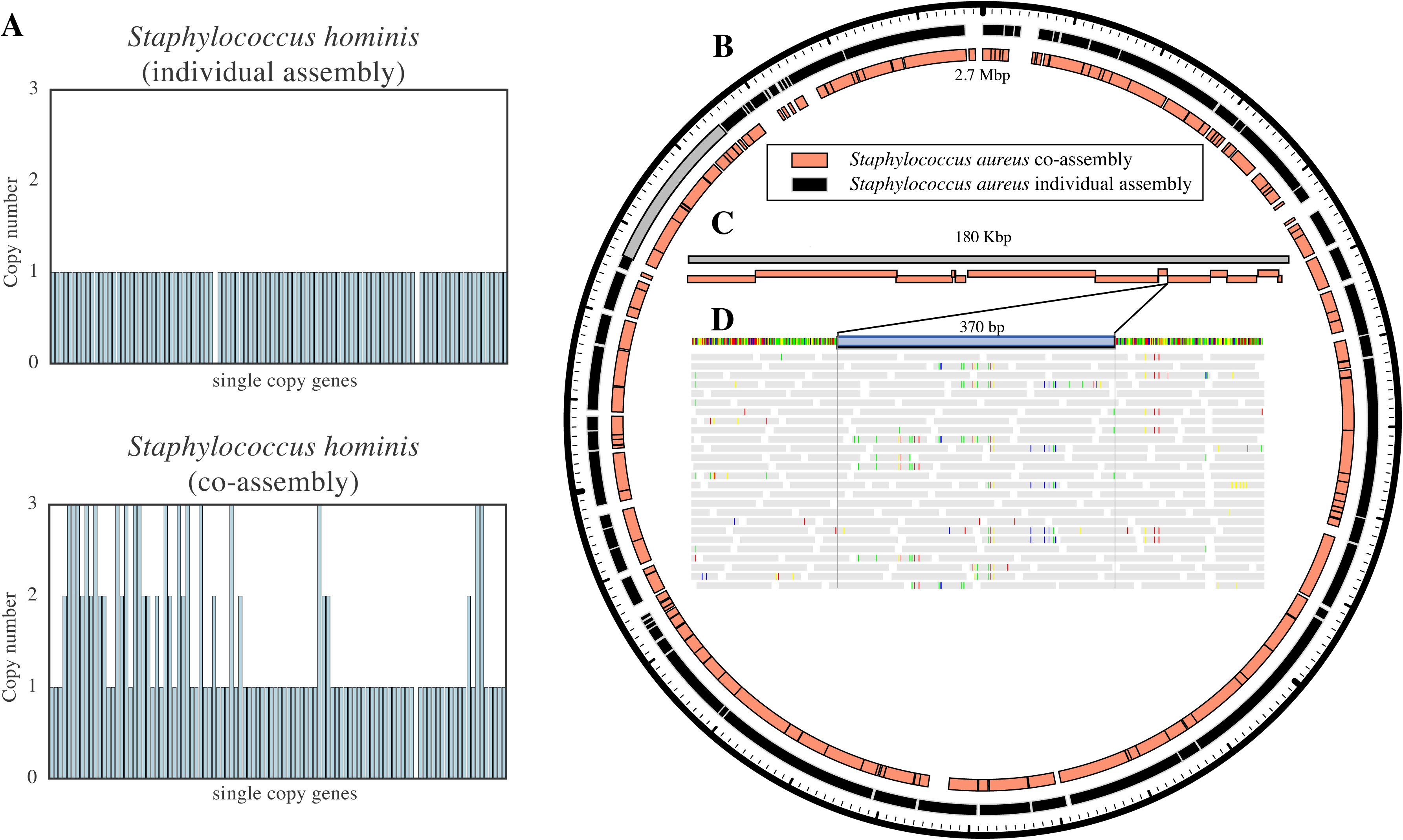
Strain heterogeneity reduces genome assembly quality and causes fragmentation in areas of extensive population-level variation. (**A**) The 104 universal bacterial single copy genes from checkM reported for a *Staphylococcus hominis* bin are shown on the x-axis, and their copy number is reported on the y-axis. Co-assembly resulted in many duplicate and triplicate single copy genes (compare top and bottom panels). (**B-D**) The *Staphylococcus aureus* bin attained from co-assembly is more fragmented than that from an individual assembly. (**B**) Scaffolds from both bins are aligned to a complete reference genome (2.7 Mbp). (**C**) Scaffolds from the co-assembly are aligned to a single scaffold (shown in grey in **B**) from the individual assembly. (**D**) Reads from all samples aligned to a gap in the alignment in (**C**). Reads mapped to the area where co-assembly failed to recover a genome sequence (highlighted in blue) show signs of population-level strain variation (**D**).

For *Staphylococcus aureus*, both co-assembly and individual assembly resulted in near-complete and uncontaminated genomes. However, alignment of the scaffolds from both *S. aureus* assemblies to a complete *S. aureus* reference genome showed that the genome from the co-assembly was more fragmented than that from the single sample assembly. (Figure 2BC). Fragmentation was also concentrated in areas of extensive population variation, as evident based on SNPs between metagenome reads and the genome sequence (Figure 2D). Genome fragmentation in sites of elevated strain variation could systematically decrease measures of within-population heterogeneity that rely on mapping reads to reconstructed genome sequences (Bendall et al., 2016; Quince et al., 2016).

It is both logical, based on the well known effects of sample complexity, and clear from the analysis of human microbiome samples presented here, that assembly of data from individual samples followed by de-replication has major advantages over co-assembly (especially as co-assembled genomes can be included in the de-replication process). Because it relies on Mash, dRep can only be used if the genomes in the comparison set are >50% complete. dRep combines checkM for completeness-based genome filtering (Parks et al., 2015), Mash (Ondov et al., 2016) for fast grouping of similar genomes, gANI (Varghese et al., 2015) or ANIm (Richter and Rosselló-Móra, 2009) for accurate genomic comparisons, and Scipy (Jones et al., 2001) for hierarchical clustering. In the case of viruses and plasmids, dRep requires use of an independent method to estimate genome completeness because there are no established metrics for this in checkM.

dRep is easy to use, highly customizable, and parallelizable. The algorithm has the sensitivity of gANI but scales more efficiently with the size of the genome set. If desired, dRep can perform rapid pairwise genomic comparisons (without de-replication) to enable visualization of the degree of similarity among groups of similar genomes (**Supp. Figure S2**). This may be particularly valuable for classifying strains as indistinguishable vs. different, a task that is important, for example, for strain tracking. For the full source-code, installation instructions, and manual, see https://github.com/MrOlm/drep.

## Acknowledgements

Funding was provided by the Sloan Foundation (http://www.sloan.org/, grant number: G 2012-10-05, PI: JFB) and the National Institutes of Health (NIH; award reference number 5R01-AI-092531). This material is based upon work supported by the National Science Foundation Graduate Research Fellowship under Grant No. DGE 1106400.

### Supplemental Information

**Supplemental Figure S1: De-replication of individual assemblies vs. co-assembly.** The methods of individual assembly and de-replication vs. co-assembly are shown for a sample series of metagenomic shotgun datasets. If there are two closely related strains changing abundance over the sample series, co-assembly will likely result in a fragmented genome. If there are some samples in which one strain or the other is dominant, individual assemblies may result in high-quality genomes. During the de-replication process the best genome from all individual assemblies is chosen, leading to better genomes compared with co-assembly alone.

**Supplemental Figure S2: Visualization of dRep primary and secondary clusters.** The first step of dRep is comparing genomes in a pair-wise manner using Mash. The resulting dendrogram (**A**) is part of the output of the program by default, and allows the user to visualize the global Mash relationship of all bins, as well as the cutoff for primary clusters (the black dotted line at 90% Mash ANI). A dendrogram for each primary cluster is also produced (**B, C**). This allows the user to visualize the secondary clustering relationship (using ANIm or gANI), as well as the cutoff for secondary clusters of “same” genomes (black line at 99% ANI). The red dotted line is the value of the lowest ANI resulting from a self-vs-self alignment of each genome in the primary cluster and represents a “limit of detection” of sorts.

**Supplemental Data 1: dRep.** A clone of the dRep program (version 0.3.3) that is available at https://github.com/MrOlm/drep.

**Supplemental Data 2: dRep manual.** A PDF reconstruction of the dRep program manual that is available at drep.readthedocs.io/en/latest/.

## References

Bendall ML, Stevens SL, Chan L-K, Malfatti S, Schwientek P, Tremblay J, et al. (2016). Genome-wide selective sweeps and gene-specific sweeps in natural bacterial populations. ISME J 10: 1589–1601.

Eren AM, Esen ÖC, Quince C, Vineis JH, Morrison HG, Sogin ML, et al. (2015). Anvi’o: an advanced analysis and visualization platform for ‘omics data. PeerJ 3: e1319.

Gibson MK, Wang B, Ahmadi S, Burnham C-AD, Tarr PI, Warner BB, et al. (2016). Developmental dynamics of the preterm infant gut microbiota and antibiotic resistome. Nat Microbiol 16024.

Graham E, Heidelberg J, Tully B. (2016). BinSanity: Unsupervised Clustering of Environmental Microbial Assemblies Using Coverage and Affinity Propagation. bioRxiv. e-pub ahead of print, doi: 10.1101/069567.

Jones E, Oliphant T, Peterson P. (2001). SciPy: Open source scientific tools for Python. URL Httpscipy Org.

Lee ST, Kahn SA, Delmont TO, Hubert NA, Morrison HG, Antonopoulos DA, et al. (2016). High-resolution tracking of microbial colonization in Fecal Microbiota Transplantation experiments via metagenome-assembled genomes. bioRxiv. e-pub ahead of print, doi: 10.1101/090993.

Luo C, Knight R, Siljander H, Knip M, Xavier RJ, Gevers D. (2015). ConStrains identifies microbial strains in metagenomic datasets. Nat Biotechnol. e-pub ahead of print, doi: 10.1038/nbt.3319.

Olm MR, Brown CT, Brooks B, Firek B, Baker R, Burstein D, et al. (2017). Identical bacterial populations colonize premature infant gut, skin, and oral microbiomes and exhibit different in situ growth rates. Genome Res gr-213256.

Ondov BD, Treangen TJ, Melsted P, Mallonee AB, Bergman NH, Koren S, et al. (2016). Mash: fast genome and metagenome distance estimation using MinHash. Genome Biol 17. e-pub ahead of print, doi: 10.1186/s13059-016-0997-x.

Parks DH, Imelfort M, Skennerton CT, Hugenholtz P, Tyson GW. (2015). CheckM: assessing the quality of microbial genomes recovered from isolates, single cells, and metagenomes. Genome Res 25: 1043–1055.

Probst AJ, Castelle CJ, Singh A, Brown CT, Anantharaman K, Sharon I, et al. (2016). Genomic resolution of a cold subsurface aquifer community provides metabolic insights for novel microbes adapted to high CO2 concentrations. Environ Microbiol n/a-n/a.

Quince C, Connelly S, Raguideau S, Alneberg J, Shin SG, Collins G, et al. (2016). De novo extraction of microbial strains from metagenomes reveals intra-species niche partitioning. http://biorxiv.org/lookup/doi/10.1101/073825 (Accessed September 12, 2016).

Raveh-Sadka T, Firek B, Sharon I, Baker R, Brown CT, Thomas BC, et al. (2016). Evidence for persistent and shared bacterial strains against a background of largely unique gut colonization in hospitalized premature infants. ISME J. e-pub ahead of print, doi: 10.1038/ismej.2016.83.

Raveh-Sadka T, Thomas BC, Singh A, Firek B, Brooks B, Castelle CJ, et al. (2015). Gut bacteria are rarely shared by co-hospitalized premature infants, regardless of necrotizing enterocolitis development Kolter R (ed). eLife 4: e05477.

Richter M, Rosselló-Móra R. (2009). Shifting the genomic gold standard for the prokaryotic species definition. Proc Natl Acad Sci 106: 19126–19131.

Sczyrba A, Hofmann P, Belmann P, Koslicki D, Janssen S, Droege J, et al. (2017). Critical Assessment of Metagenome Interpretation – a benchmark of computational metagenomics software. bioRxiv. e-pub ahead of print, doi: 10.1101/099127.

Sharon I, Morowitz MJ, Thomas BC, Costello EK, Relman DA, Banfield JF. (2013). Time series community genomics analysis reveals rapid shifts in bacterial species, strains, and phage during infant gut colonization. Genome Res 23: 111–120.

Tyson GW, Chapman J, Hugenholtz P, Allen EE, Ram RJ, Richardson PM, et al. (2004). Community structure and metabolism through reconstruction of microbial genomes from the environment. Nature 428: 37–43.

Varghese NJ, Mukherjee S, Ivanova N, Konstantinidis KT, Mavrommatis K, Kyrpides NC, et al. (2015). Microbial species delineation using whole genome sequences. Nucleic Acids Res 43: 6761–6771.

Vineis JH, Ringus DL, Morrison HG, Delmont TO, Dalal S, Raffals LH, et al. (2016). Patient-Specific *Bacteroides* Genome Variants in Pouchitis. mBio 7: e01713–16.

